# The effect of mentee and mentor gender on scientific productivity of applicants for NIH training fellowships

**DOI:** 10.1101/2021.02.02.429450

**Authors:** Hao Yu, Kristine A. Willis, Aviva Litovitz, Robert M. Harriman, Matthew T. Davis, Payam Meyer, Brad Busse, Rebecca A. Meseroll, Hashanthi D. Wijayatilake, Matthew J. Perkins, James M. Anderson, George M. Santangelo

## Abstract

Several studies have suggested that women in science are less productive than men, and that this gap contributes to their under-representation in the ranks of senior researchers. However, few studies have examined the role of mentoring, and in particular mentor gender, on the productivity of female scientists early in their careers. Such efforts are limited by the difficulties of unambiguously linking mentees to their mentors and measuring the research productivity resulting from those relationships. Here we use our novel author disambiguation solution to investigate the role of self-identified gender in mentorship of 12,932 trainees who either successfully or unsuccessfully applied to the National Institutes of Health for research fellowships between fiscal years 2011 and 2017, applying a multi-dimensional framework to assess productivity. We found that, after normalizing for the funding level of mentors, the productivity of female and male mentees is indistinguishable; it is also independent of the gender of the mentor, other than in measures of clinical impact, where women mentored by women outperform other mentee-mentor dyads.

## Introduction

It is well established that women who pursue careers in biomedical sciences face formidable barriers. Gender bias may contribute, since for example university professors given identical resumes headed by either a male or female name are more likely to view male lab technician candidates as competent and hirable, and to offer male candidates higher salaries [1]. After controlling for prior productivity and achievements, reviewers of postdoctoral fellowship applications view male candidates as more meritorious [2, 3]. Women receive harsher teaching evaluations, are less likely to be judged as stars in their field by reviewers of R01 applications to the National Institutes of Health (NIH), and after normalizing for a number of confounding factors, including journal of publication, number of authors, and seniority, their work accrues fewer citations than that of their male colleagues [4–7]. Though women in the life sciences represent only slightly more than a third of all tenured or tenure-track professors employed by universities or four-year colleges, they are awarded roughly half of all doctoral degrees [8]. Importantly, women of color face a double bind that hinders their entry into, and retention and advancement in, biomedical careers [9].

While the overall progress and remaining challenges experienced by women in biomedicine have been widely discussed, the potential effects of mentorship on their career progress have received relatively little attention. Recently though, a small number of papers have raised the possibility that mentorship, career progress, and gender interact in important ways. Among the very small and elite group of science faculty who have funding from the Howard Hughes Medical Institute, have been inducted into the National Academy of Sciences, and/or have won a Nobel prize, men are significantly more likely to employ other men as postdoctoral fellows; members of the National Academy of Science, which is 85% male, train 58% of future faculty [10]. In contrast with these data, which suggest that the careers of female mentees may be disadvantaged by their exclusion from elite male networks, other work suggests that having a female mentor is an advantage to mentees; for example, a study of roughly 900 PhD students at a single university found that on average, doctoral candidates studying biology under female advisors publish approximately 10% more papers, and tend to publish in more influential venues, than those with male advisors [11]. In the adjacent field of chemistry, a larger study found that women who chose a female advisor for their doctoral studies were more productive and more likely to go on to faculty positions than those who chose a male advisor [12]; however, this work systematically excluded students with Chinese and Korean names because of the difficulty in assigning them algorithmically to a gender, significantly weakening its conclusions.

These inconsistent findings suggest that a more comprehensive analysis of mentorship and gender might identify factors that either exacerbate or mitigate the barriers faced by women in science. In addition to their small sample sizes, previous studies have been further limited by inaccuracy of assigning mentees to their mentors, difficulty verifying the gender of both, and/or limiting the measurement of research productivity such as publications to the training experience. To overcome these drawbacks, we studied mentee-mentor relationships among applicants for individual training fellowships from the NIH. Most applicants for NIH fellowships choose to self-identify gender, and all are required to identify their mentor(s). Almost all mentors are also NIH-funded investigators who self-identify gender in their own applications. We used our novel disambiguation method to accurately document mentee research productivity. Since NIH fellowships cover topics ranging from computational biology, synthetic chemistry, and biophysics, to epidemiology and clinical psychology, this allowed the construction of a large, reliable set of self-reported mentee-mentor pairs spanning a wide range of scientific disciplines, which we analyze here.

## Results

We began our analysis with 18,600 applications for individual fellowships (predoctoral mechanisms F30 and F31, and postdoctoral mechanism F32, K01, K08, K23, and K99) submitted to NIH between fiscal years (FY) 2011 and 2017. Over this time frame, women and men applied for fellowships in similar numbers and received awards at the same rate; this is true if applications for pre- or post-doctoral fellowships are considered either together (two leftmost bars, **Figure 1a**) or separately (two leftmost bars, Figure **1b** and two leftmost bars, **Figure 1c**). As mentioned above, a unique feature of this dataset is that applicants communicated to NIH the names of the independent investigators who would act as their sponsor(s) or mentor(s). NIH requires that any independent investigator named by a fellowship applicant in either of these capacities (for simplicity, referred to hereafter as a ‘mentor’) must demonstrate an understanding of the candidate’s training needs, as well as the ability and commitment to assist in meeting these needs. Mentors must provide a letter of support as a part of the fellowship application package, and an evaluation of this statement, as well as evidence of successful outcomes for the mentor’s past mentees, are among the explicit criteria that review panels are instructed to use in their evaluation. Focusing on mentors identified in fellowship applications allowed us to analyze the relative impact of gender on mentee productivity, though applicants may have access to other individuals who provide advice and guidance.

**Figure 1.**
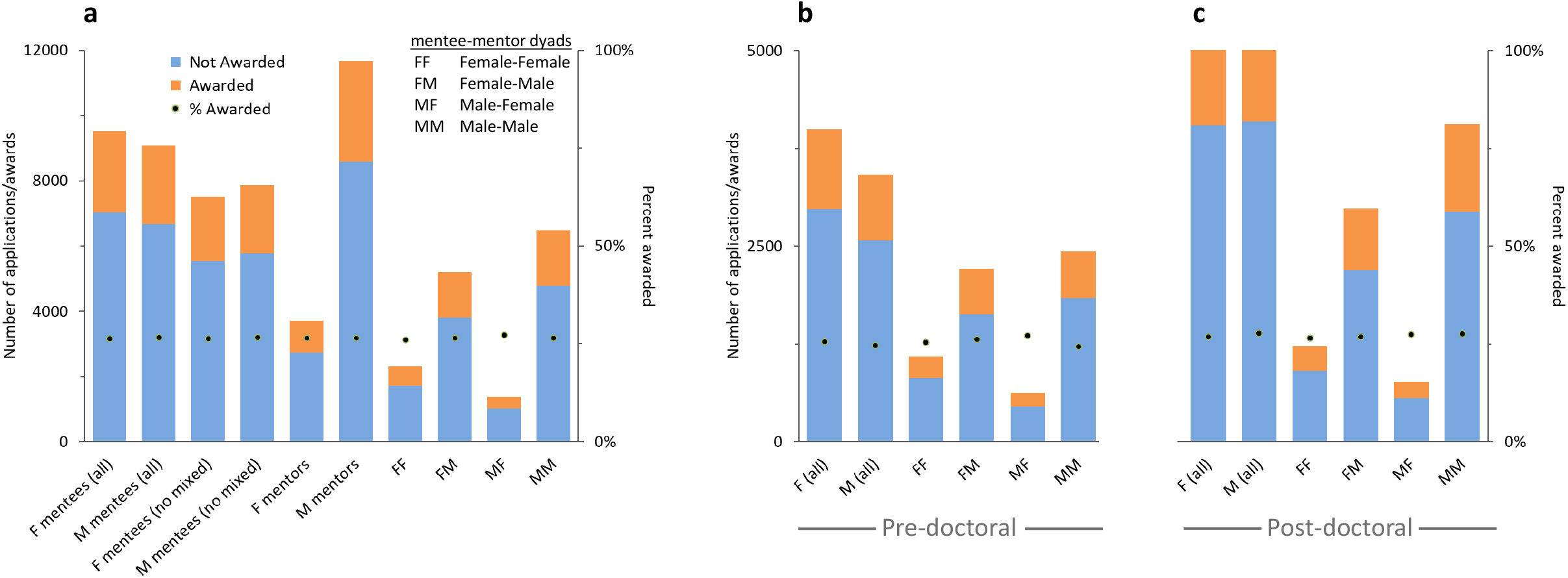
Fellowship applications and awards by mentee and mentor gender. NIH fellowship applications (pre-doctoral: F30 and F31; post-doctoral: F32, K01, K08, K23, and K99) submitted by females (F) and males (M) in FY11 through FY17. Light blue and orange bars represent unawarded and awarded applications, respectively; black dots indicate award rates (secondary Y axis). **(a)** The first two bars represent applications from all mentees with single-gender or mixed-gender mentors; the third and fourth bars represent applications from mentees with single-gender mentors, separated by mentee gender; the fifth and sixth bars represent applications from mentees with single-gender mentors, separated by mentor gender. The remaining four bars show the data analyzed by mentee-mentor dyads for mentees with single-gender mentors. Mentee gender is presented first, mentor gender second, e.g., FM = female mentees with male mentors. **(b and c)** Same data as in (a), analyzed by pre- or post-doctoral career stage of the mentee applicant, respectively. The first two bars represent applications from all mentees with single-gender or mixed-gender mentors. The remaining four bars show the data analyzed by mentee-mentor dyads for mentees with single-gender mentors. There is no statistically significant difference in award rate for any group.

Consistent with previously published data on the proportion of women in the R01 applicant pool [13], approximately 30% of mentors in our dataset are female. A subset of applications (17%) were submitted by mentees who, either in a single application or in two or more different applications, identified both male and female independent investigators as mentors. To simplify our analysis and avoid double counting, we removed those applications, which reduced the proportion of female mentors from 30.4% to 25.6% but had no effect on mentee award rates (third and fourth bars, **Figure 1a**). Mentees of either gender who list exclusively female mentors, and those who list exclusively male mentors, have identical award rates (fifth and sixth bars, **Figure 1a**); further dividing applicants into four dyads based on the gender of both mentee and mentor also fails to identify any gender-based differences in award rates, regardless of whether pre- and post-doctoral fellowships are considered together (last four bars, **Figure 1a**) or separately (last four bars, **Figure 1b, c**).

We next asked if the genders of the mentee/mentor dyads influenced mentee research productivity. Most analyses of productivity are limited to awardees due to the need to rely on the grant number cited in the resulting publications to link an investigator to his or her papers. One major drawback of this method is that it is unable to assign papers to unsuccessful applicants. Past studies have attempted to address this problem by creating a restrictive set of criteria to match grant applicants with the papers they have authored, such as requiring identical first names, last names, and institutional affiliations. However, such methods will fail to match authors who change names or institutions, or where mismatches have been introduced as the result of typos, inconsistent spellings, or the inconsistent use of a middle initial. To address this problem, we developed a disambiguation solution that used article-level metadata [14–16] to assign 24.5M unique papers from the PubMed database to 16.0M unique author names, then used a novel neural network model trained on ORCID identifiers to determine whether author-publication pairs refer to variant representations of the same person (see **Methods** for details). For example, our model (**Figure 2**) can determine whether hypothetical records listing Jane Smith and Jane M. Smith were the same person, or two different people, based on variables that include institutional affiliation, co-authorship, and article-affiliated Medical Subject Heading (MeSH) terms. We then matched unambiguously identified author and applicant names. Importantly, the model does not require last names to match, so women who change their name can be successfully merged. We used this method to reduce the 16.0M unique author names to 13.3M disambiguated people; the F1 score for people with at least one NIH application is 0.945, indicating both high precision and high recall. Disambiguation of the people associated with the fellowship applications in our dataset indicates that we have captured a large fraction of NIH trainees and their mentors, since the papers of these unambiguously identified applicants together amount to 57.7% of all publications since 2011 that cite NIH grant support.

**Figure 2.**
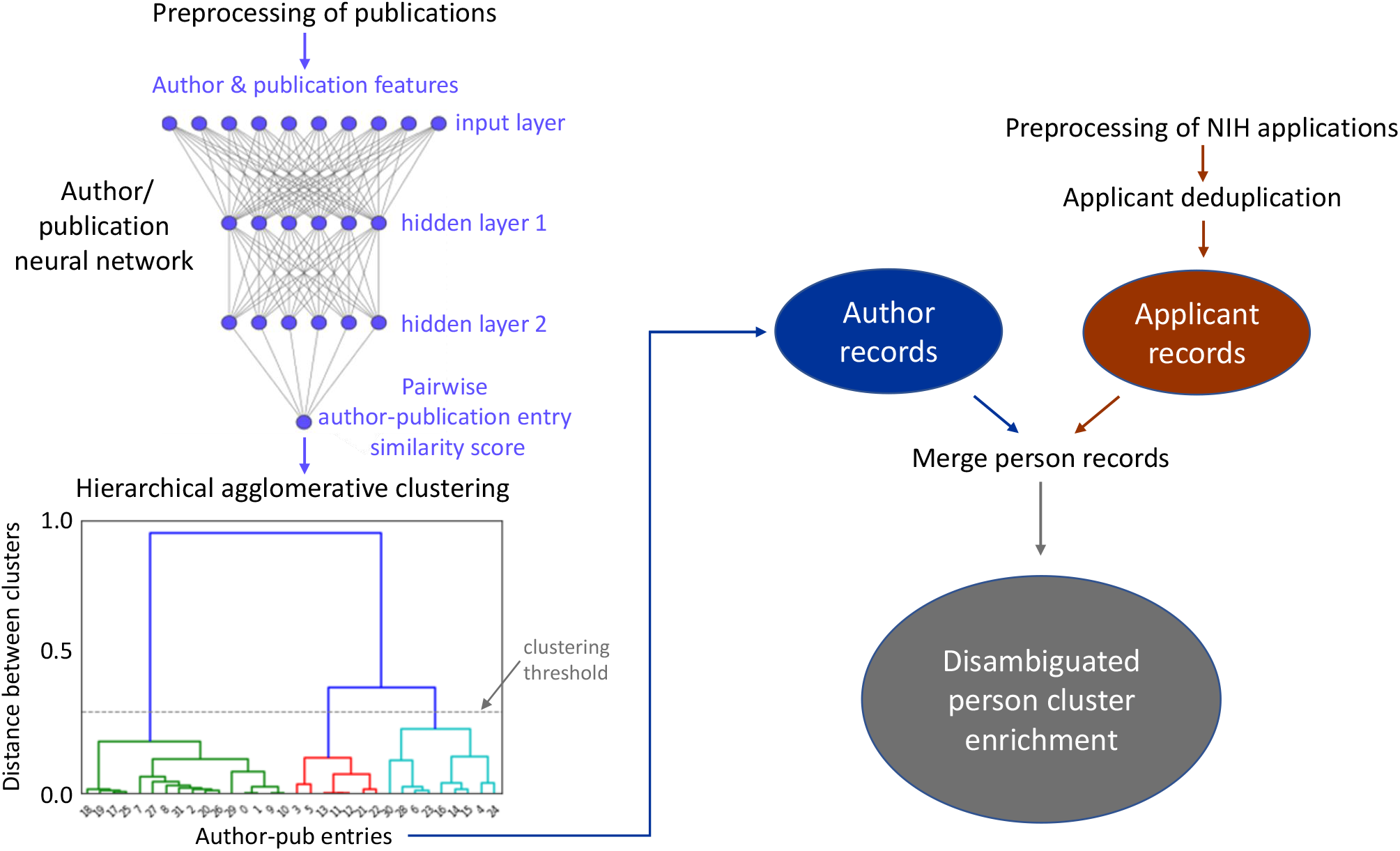
Disambiguating authors and NIH applicants. Graphical representation of the workflow used to disambiguate unique author names and link publications and NIH applications to specific authors and applicants. The process began by assigning 24,453,076 unique publications to 15,985,142 unique author names, and resulted in 13,324,796 disambiguated people. A fully connected neural network with two hidden layers, trained on a series of author and publication features, generated pairwise author-publication entry similarity scores (left side of the illustration; see Methods). Those similarity scores were used by a hierarchical agglomerative clustering algorithm to merge author-publication entries, resulting in disambiguated PubMed author records (blue oval). In parallel, preprocessing of NIH applications and applicant deduplication (see Methods) generated applicant records (red oval). Subsequent matching and merging of disambiguated PubMed author records with deduplicated applicant records generated disambiguated author profiles that contain specific linkages to a person’s publications and NIH applications. The disambiguated person records were enriched with the person’s metadata and data for each publication and grant application to facilitate downstream analyses (grey oval).

Our data show that male-male dyads, and more specifically, male-male post-doctoral dyads, have more publications prior to the time of their first application (set at time = 0, **Figure 3a-c**). Male-male post-doctoral dyads also start out with more papers in the top decile of Relative Citation Ratio (RCR) values (**Figure 3d-f**); RCR is an article-level, field- and time-normalized measure of scholarly influence [17]. This early advantage in publication and citation metrics is maintained for years after the time of first application (**Figure 3a-f**), consistent with a hysteretic process. Beginning around the time of application, the number of highly influential (top decile) papers authored by female-female post-doctoral dyads begins to diverge from the number published by male-female and female-male dyads (**Figure 3e**). Eight years after their first application, this difference is outside the confidence interval; it should be noted that this is roughly correlated with the point at which women are more likely to leave academia [18]. However, median RCR values are indistinguishable for all four types of mentee-mentor dyads over the entire eighteen-year time frame of our analysis (**Figure 3g-i**), and applicants for pre-doctoral fellowships exhibit no meaningful differences outside the 95% confidence interval for any of these productivity measures, regardless of mentee or mentor gender, among applicants for pre-doctoral fellowships (**Figure 3c, f, i**).

**Figure 3.**
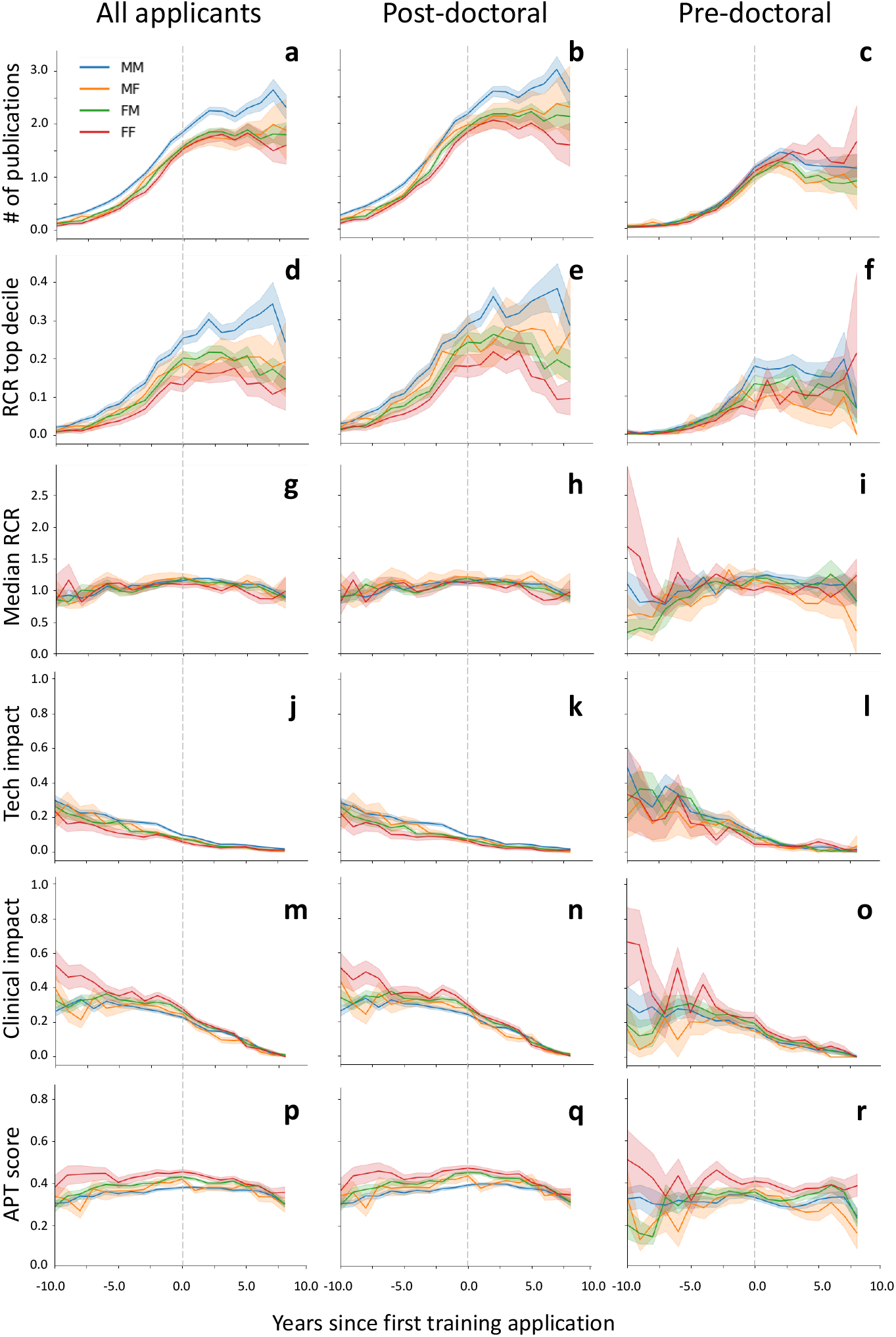
Mentee productivity over time. Six different measures of mentee research productivity, shown per mentee per year, where the first fellowship application is set to time=0 (vertical dashed grey line): **a-c**, mean number of publications; **d-f**, mean number of high-influence publications (defined as having an Relative Citation Ratio (RCR) value in the top decile); **g-i**, median RCR; **j-l**, technological impact; **m-o**, clinical impact; **p-r**, mean Approximate Potential to Translate (APT) score (see Methods for additional details). First column, all (both post- and pre-doctoral fellowship applicants; second column, post-doctoral applicants only; third column, pre-doctoral applicants only). Shaded regions indicate 95% confidence intervals, determined via bootstrap analysis. The dyads are annotated with mentee gender first, mentor gender second (e.g., FM – female mentees with male mentors)

Publication and citation metrics are the typical, but not the only, measure of scholarly contribution to scientific progress [19]. Biomedical research also leads to patentable inventions/technological (tech) impact, measured by the citation of publications by patents, and clinical impact, measured by the citations of publications by clinical trials and guidelines. Female-female dyads appear to have less tech impact (**Figure 3j-l**) and more clinical impact (**Figure 3m-o**), in both pre- and post-doctoral applicant populations; since these forms of citations are slower to accrue than citations to peer-reviewed publications, censoring (the absence of hypothetical future citations; [20]) makes it difficult to determine whether these differences are maintained. However, clinical impact can also be measured with APT (Approximate Potential to Translate) scores, which are machine-learning based predictions of future clinical citations [21]; these predictions are particularly useful because they are less subject to censoring. Both before and after applying for a fellowship, APT scores are highest for female-female dyads. Together, the greater number of clinical citations (**Figure 3m-o**) and higher APT scores (**Figure 3p-r**) indicate that this dyad generates the highest level of clinical impact.

Although small dollar amounts may sometimes be budgeted for a training course or similar expense, NIH fellowships generally provide salary only. The productivity of the mentee is therefore heavily reliant on the amount of research funds available to the mentor. Interestingly, for each of the four mentee-mentor dyad categories, mentors of post-doctoral fellowship applicants have a higher level of median total costs, adjusted for inflation to 2019 dollars by using the Biomedical Research and Development Price Index (BRDPI; **Figure 4**). This is unlikely to be explained by the institutional affiliation of mentors, since pre-doctoral and post-doctoral applications distribute similarly across institutions receiving widely different levels of NIH support, regardless of whether aggregate inflation-adjusted funding or dollars per investigator are considered (see **Supplemental Data**). Our data are also consistent with previously published work [13] showing that on average, women hold fewer awards (**Supplemental Data**) and have fewer research dollars than men (**Figure 4**). This disadvantage does not influence the chance of winning a fellowship award for applicants with female mentors (**Figure 1a**), but might have an impact on mentee productivity. We therefore normalized the number of publications, scholarly influence, tech impact, and clinical impact of mentees to the total amount of NIH funding held by their mentors.

**Figure 4.**
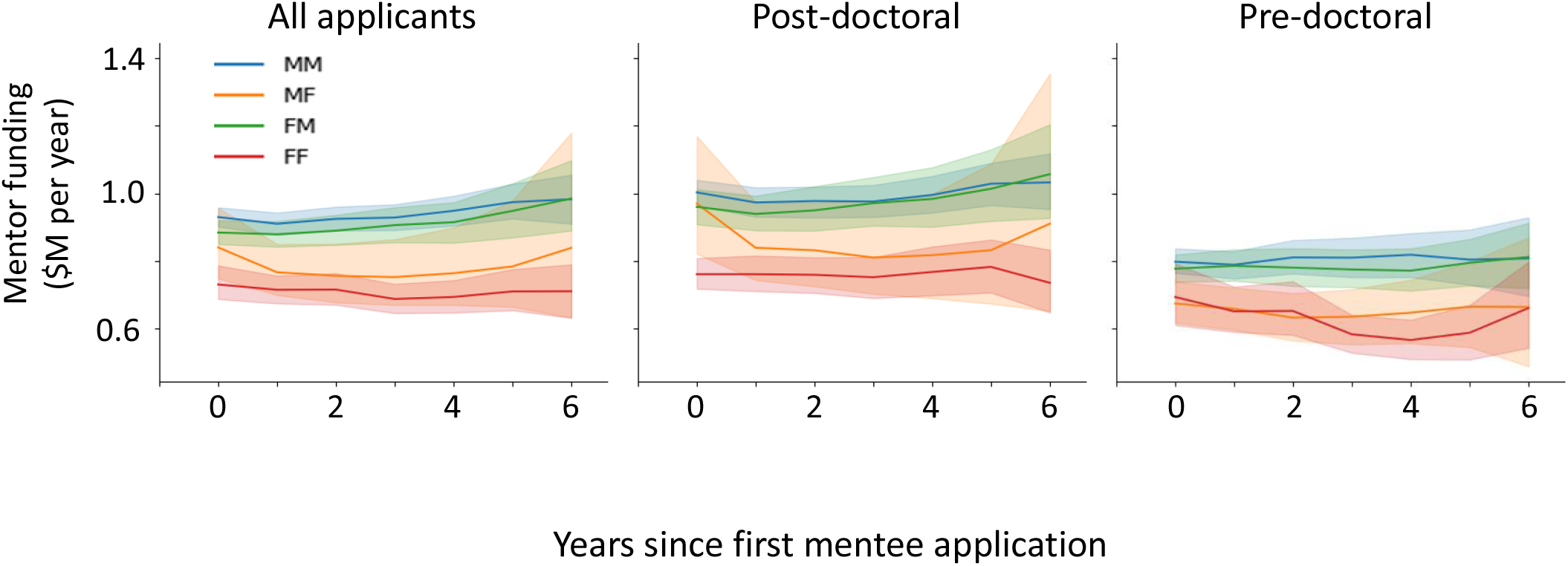
Mentor funding over time. Mentor funding levels per mentee for gender-based mentee-mentor dyads in the six years following a mentee’s first fellowship application (set to time=0). Left graph, all (both post- and pre-doctoral fellowship applicants; middle graph, post-doctoral applicants only; right graph, pre-doctoral applicants only). Shaded regions indicate 95% confidence intervals, determined via bootstrap analysis. The dyads are annotated with mentee gender first, mentor gender second (e.g., FM – female mentees with male mentors)

Strikingly, adjusting for the funding available to mentors eliminates the advantage of male-male post-doctoral dyads in number of publications (**Figure 5a, b**), scholarly influence (**Figure 5d, e**), and tech impact (**Figure 5j, k**). More than simply closing the gap, after adjusting for funding, female-female post-doctoral dyads have a slightly higher number of papers immediately prior to the time of their first application for a fellowship (**Figure 5b**). They also have greater clinical impact (**Figure 5m, n**) and higher APT scores (**Figure 5p, q**). Female-male dyads also have higher APT scores than male-female or male-male dyads (**Figure 5q**), suggesting that female mentees in general are more successful at producing clinical impact. Finally, the median RCR values of post-doctoral dyads are largely indistinguishable (**Figure 5g, h**), and the high signal to noise ratio for the four pre-doctoral dyads could in part be responsible for the failure to detect differences in those funding-adjusted metrics (**Figure 5c, f, I, l, o, r**).

**Figure. 5.**
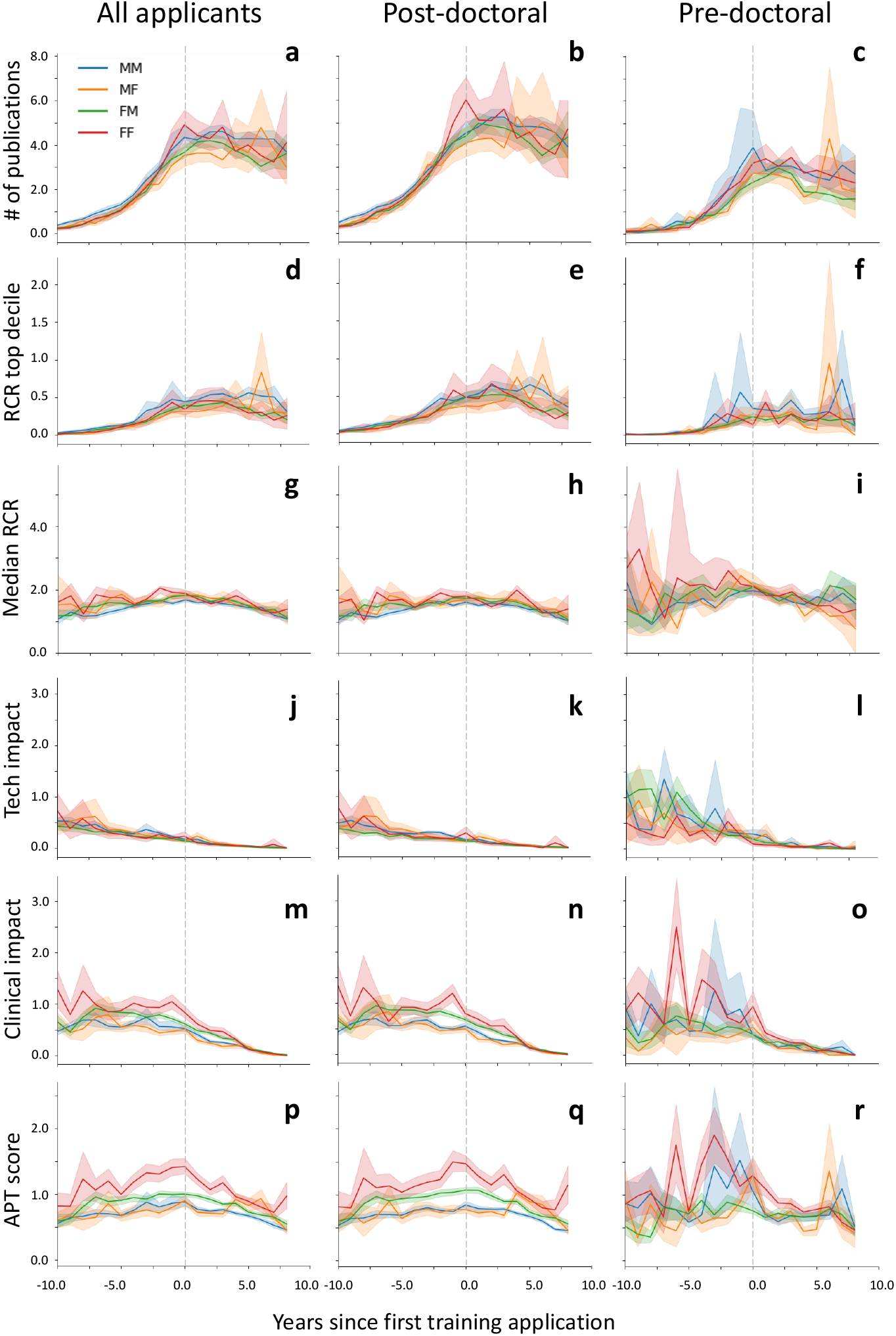
Funding-normalized mentee productivity over time. In order to account for the effect of differing mentor resources on mentee productivity, Figure 3 data was re-analyzed to normalize for mentor funding levels. As in Figure 3, the six different measures of mentee research productivity are presented for gender-based mentee/mentor dyads, normalized per million mentor Principal Investigator dollars: Mean number of publications (**a-c**), mean number of high-influence publications (**d-f**), median RCR of all publications (**g-i**), technological impact (**j-l**), clinical impact (**m-o**), and APT score (**p-r**). Data are analyzed per mentee per year, grouped by post- or pre-doctoral career stage of the mentee applicant (columns left to right), and are presented across a time scale of ten years before and after a mentee’s first post-doctoral (middle column) or pre-doctoral (right column) fellowship application. The first fellowship application is indicated at time=0 (vertical dashed grey line). Shaded regions indicate 95% confidence intervals, determined via bootstrap analysis. When a mentee is linked to multiple mentors, their funds are averaged. See Methods for more details. The dyads are annotated with mentee gender first, mentor gender second (e.g., FM – female mentees with male mentors)

## Discussion

Previous reports have found that two prominent and potentially interrelated barriers faced by female scientists are the postdoc-to-faculty transition to independence and the greater difficulty in achieving higher levels of influence via citation of their research outputs. Our analysis of the productivity to date of FY11-FY17 applicants for individual NIH fellowships confirmed the latter observation: female mentees produce fewer papers in the top decile of RCR values, as well as fewer papers overall. However, normalizing to mentor funding levels eliminates both of those gaps. We also found that the median RCR for female and male mentees are indistinguishable, and male mentees, especially when they are paired with male mentors, have lower clinical impact. Taken together, the data indicate that, if there is any gender-based difference in mentorship at all, it manifests as an advantage of female mentoring of female mentees in producing clinically relevant research.

Our data suggest that the initial appearance of lower productivity of female mentees might be a direct result of the funding gap between independent female and male investigators that has already been noted in the literature [13]. In addition to confirming that gap, we found that pre-doctoral mentors in each of the four dyad categories are less well funded than post-doctoral mentors (**Figure 4**). Since institutional affiliation makes at most a minor contribution to this difference, the funding gap between pre- and post-doctoral mentors may indicate a preference on the part of individual independent investigators. If so, less well-funded scientists are more willing or able to sponsor individual pre-doctoral applications, while well-funded investigators prefer to sponsor post-doctoral fellowships.

Among the benefits of data-driven decision making is the potential to inform the pursuit of desired goals and to dispel misconceptions that might result in unhelpful guidance or policy decisions. Large-scale metascience uses many different types of information towards this end (e.g. grant applications, publications, patents, clinical trials), and requires the careful generation of accurate linkages among these disparate sources of data. Author names, applicant names, affiliations and other metadata are presented in many different variations (e.g. John D Smith vs. JD Smith vs. John Smith); identification of a single person, accurately linked to their full publication and application record, requires a rigorous method of name disambiguation. Without such methodology in hand, attempts at meaningful analysis are plagued by incomplete records and/or multi-counting errors. Our newly developed high-performance Artificial Intelligence/Machine Learning (AI/ML)-based disambiguation method improves on previous attempts [22–24] to address this problem, achieving an F1 score of 0.945 and allowing us to clean the large datasets of PubMed author names and NIH grant applicants then integrate that information to create the dataset used in this study.

Our disambiguation solution also allows us to overcome several challenges created by the common practice of linking outputs to funded grants. As we have shown here, it allows publications to be linked to applicants who have not yet (or ever) received an NIH award. It also allows grants to be linked to papers if the authors failed to cite their award or cited it in a non-specific way, such as acknowledging support from the NIH without providing an identifying number. Finally, person-level links solve the problem posed by publications in journals that lack an acknowledgements section.

We have relied on these person-level links, and the availability of self-identified information provided by NIH training fellowship applicants, to investigate the role of mentee and mentor gender on mentee productivity. This has removed the inevitable errors associated with a reliance on tenuous assumptions in defining mentee-mentor relationships. Of course, not all mentoring occurs in the context of a formal relationship with one or more doctoral or post-doctoral advisors. A variety of scenarios, ranging from structured, regular meetings with thesis committee members to informal, transient interactions in which a more senior scientist gives technical or career advice to a junior colleague, may be interpreted as mentorship. While often critically important, methods capable of fully capturing these networks must go beyond a simple analysis of co-authorship [25].

By identifying the outsized clinical impact of female mentees, which is a previously unappreciated contribution to biomedicine, our analysis demonstrates the value of using a multifaceted framework for measuring research productivity. The development of additional metrics that measure other factors supporting scientific progress, such as rigor/reproducibility and data sharing, should further improve analyses that can be effective in informing and guiding policy [19]. We are also now poised to go beyond the current analysis to investigate the role of specific mentor and mentee characteristics (e.g. career stage, mentoring track record, affiliation, previous publication record) in promoting the successful transition to productive independent careers. This type of information has the potential to inform guidance and policy-making that provide robust support for the scientific enterprise, as former mentees in turn train the next generation of scientists and perpetuate the cycle of progress that has now proved its worth as a means of advancing knowledge that improves human health.

## Methods

### Author and applicant name disambiguation methodology

Author name disambiguation was carried out in two stages (**Figure 2**). The first stage was disambiguation of PubMed authors and deduplication of grant applicants. The second stage of the process involved matching and merging the disambiguated PubMed author records with the deduplicated applicant records to generate disambiguated author profiles that contain specific linkages to a person’s publications and NIH funding.

#### PubMed author disambiguation

PubMed author disambiguation was performed in a similar fashion as described previously [14–16]. Essentially, authors were disambiguated in the same first initial last name (FILN) block using hierarchical agglomerative clustering algorithm based on pairwise similarity.

During preprocessing, each individual author on a publication was listed separately to form author-publication entries. Author and publication metadata (author name, author affiliation, coauthor names, location, journal name, linked grant numbers, clinical trials, and patents), citations, and content features (MeSH keywords, title tokens, broad subject terms) were collected or extracted and stored in author-publication entries. The author-publication entries were then grouped based on the author FILN.

In order to cluster the author-publication entries within each FILN block to form disambiguated author records, a fully connected neural network with two hidden layers was trained as a binary classifier to determine if two author-publication entries belong to the same author using the attributes stored in author-publication entries. The probability output of the model was considered to reflect the similarity of the input author-publication entries. Training and test datasets were generated using ORCID profiles. Hierarchical agglomerative clustering algorithm [26] was used to cluster the author-publication entries using similarity scores generated by the trained neural network model. The resulting disambiguated author records were used for the person record merging stage below.

#### Grant applicant deduplication

In parallel to PubMed author disambiguation, NIH grant applicants were deduplicated by deduplicating their Principal Investigator IDs (PIIDs). Ideally, PIIDs should map to applicants in a one-to-one relationship. However, we estimated that 10-15% of all PIIDs were duplicates. These PIIDs and their associated applications needed to be merged before linking them to the disambiguated PubMed authors. PIID deduplication was performed as following: In a preprocessing step, applications were unwound on all PIs listed on the application. The following information was then extracted: PIID, PubMed IDs (PMIDs) linked from the NIH Scientific Publication Information Retrieval and Evaluation System (SPIRES; we collected links of match case 3, 4, and 5 and then further screened them with our name matching algorithm), PMIDs resolved from grant applicants’ biosketches, and metadata such as applicant name and grant number. The unwound applications and all the extracted data associated with the application were aggregated to PIIDs.

In the deduplication step, PIIDs were combined if they met one of two criteria: 1) Consecutive PIIDs with matched applicant names; 2) Same grant number with matched applicant names. A few hundred PIIDs were also manually curated and used for deduplication. The resulting applicant records were used for the person record merging stage below.

#### Person record merging

The disambiguated PubMed authors and deduplicated grant applicants from the first stages were merged based on name matching and publication overlap for records that shared the same FILN. However, we observed that a small percentage of applicants had name variants with different FILNs caused by various reasons. The most common reasons include typos in the last names, re-arranged first/middle names, and name change due to marriage. As a result, their publications were disambiguated in different FILN blocks. To allow combining over-split records of the same applicant from different FILN blocks, we merged records across FILN if they were associated with the same PIID.

#### Enrichment

To facilitate downstream analyses, disambiguated person records were enriched by generating the best name for the person using all the name variants that appeared in their publications and grant applications, and populating any useful data for each publication and grant application in the disambiguated record.

#### Assessing performance of the disambiguation method

We developed an evaluation method using ORCID profiles which we treated as ground truth. ORCID authors were mapped to disambiguated author records by name matching and publication overlap. Precision and recall were computed accordingly. If one ORCID author was mapped to more than one disambiguated author records, the record that had the highest F1 was designated as the disambiguated author for that ORCID author. Because ORCID data are largely incomplete in terms of their publication records, we only considered PMIDs that could be found or resolved in ORCID profiles for precision calculation.

Since only about 1.8% of all disambiguated authors are associated with NIH grants and this study concerns grant applicants, we evaluated the performance of our disambiguation process in two groups of authors: those with grants and those without grants. Micro-precision, micro-recall, and micro-F1 were computed for random samples of these two groups. For one experiment, 7000-7500 samples for authors without grants and 450-500 samples for authors with grants were randomly selected to compute the performance metrics for each group. This experiment was repeated five times and the metrics were compared using unpaired Student’s t-test.

We found no statistically significant difference between the precision of disambiguated authors with grants and without grants (0.985 ± 0.003 for authors without grants vs. 0.984 ±0.007 for authors with grants, p = 0.86). In contrast, disambiguated authors with grants had significantly higher recall than disambiguated authors without grants (0.783±0.008 for authors without grants vs. 0.908±0.016 for authors with grants, p < 0.0001). F1, as a result, showed the same trend as recall (0.872 ±0.006 for authors without grants vs 0.945 ±0.010 for authors with grants, p < 0.0001).

### Identifying fellowship applications

Mentees were defined as NIH grant applicants who had applied for pre-doctoral or post-doctoral training fellowships (F30 and F31 for pre-doctoral; F32, K01, K08, K23, and K99 for post-doctoral) in fiscal years 2011-2017. 2011 was chosen as the initial analysis year since it was the first year fellowship applicants were able to self-identify their mentors/sponsors as a “Key Person” during the application process. This official record of self-identified mentee-mentor relationships was considered critical in identifying valid and accurate mentee/mentor dyads. 2017 was chosen as the final year of analysis to give the mentees sufficient time to produce publications after their fellowship application, while allowing time for reliable publication-related metrics to subsequently accrue.

A total of 57,425 applications from 37,918 mentee applicants were first identified. The following selection criteria was then applied to the dataset: 1. mentors should have a PIID in the Key Person field or the name/organization search should return a single profile match (for mentors that did not have a PIID associated with the mentee application, mentor PIIDs were added from the matched profile; n=35,999 applications from 21,856 applicants were excluded); 2. mentors should have only one PIID (those with zero or >1 were excluded; n=3,902 applications from 2,853 applicants); 3) mentees should have no more than one PIID (those with >1 PIIDs were excluded; n=414 applications and 277 applicants).

The selection criteria above yielded a dataset of 18,600 unique applications from 12,932 mentee applicants. All these mentees were initially analyzed, including those who had mixed-gender mentors (Figure 1a). To simplify the analysis, avoid double counting, and avoid conflating effects of mixed (both gender) mentors, 3,215 applications (17%) from 1,858 mentee applicants who had a combination of female and male mentors were eliminated for the subsequent analysis. This yielded a final dataset of n=15,386 applications from 11,074 mentee applicants with single-gender (i.e. female-only or male-only) mentors.

Fellowship award rates are presented as a percentage of total awards.

### Assignment of gender

Gender is self-identified by NIH applicants during the application process. Of the mentee/mentor dyads in our dataset, 94.1% of mentees and 90.09% of mentors had self-identified records of gender. Genderize (genderize.io) was used to assign gender to those without self-identified gender information, as done in previous studies [12, 27, 28]. The agreement between self-identified gender and Genderize was 98%, allowing confidence in using Genderize to populate the small portion of non-self-identified records.

Statistical significance was calculated using Fisher Tests compared to the rest of the population.

### Analysis of mentee productivity over time

Productivity metrics were analyzed for each mentee, and time-shifted for each mentee-mentor dyad such that year 0 represented the year of the mentee’s first fellowship grant application (regardless of awarded status). Mentees were split between pre-doctoral mentees who received only F30 or F31 awards over the course of the analysis period (right columns in Figures 3 and 5), and post-doctoral mentees who received any other type of award listed in the section above (middle columns in Figures 3 and 5). n=138 mentees had both pre- and post-doctoral fellowship applications and were counted in the post-doctoral group.

In each case four subpopulations of mentee/mentor dyads were measured independently, based on their respective gender: male mentee/male mentor (MM), male mentee/female mentor (MF), female mentee/male mentor (FM), female mentee/female mentor (FF). As noted above, mentees with multiple mentors of both genders were excluded from the analysis. Shaded regions indicate 95% confidence intervals, determined through bootstrap analysis.

Six research productivity metrics were examined, per mentee, per year, before and after the first fellowship application (**Figures 3 and 5**): Mean number of publications, mean number of high-influence publications (defined as having an RCR [17] in the top 10% of all NIH publications), median RCR of all publications, technological impact (fraction of publications which have been cited by at least one US patent submission), clinical impact (fraction of publications which have been cited by at least one clinical trial or guideline), Approximate Potential to Translate (APT, [21]) score (predicted clinical impact fraction based on citation trends). Dyads with no mentee publications (n=1,999, or 11.0%) were excluded from the analysis. Further, to facilitate comparisons with funding information (see below), dyads with no funding information (e.g. those with mentors entirely funded outside of NIH) were also excluded from the normalized analysis (n=3,236, or 20.0%).

The same six mentee productivity metrics noted above were subsequently re-analyzed to normalize to mentor funding levels (**Figure 5**). Mentor funding levels were defined as the amount of NIH mentor funding (averaged between mentors if multiple are linked to the same mentee) in the fiscal year of the mentee’s training application, adjusted for inflation to BRDPI 2019 dollars. Time points before the first training application use the first application’s value, and time points after use the most recent application. Each mentee’s productivity metric was divided by their mentors’ funding level, and normalized to productivity per million PI dollars. For example, mean publications per year presented in Figure 3 are presented as mean publications per million dollars per year in Figure 5.

## Notes

### Competing Interest Statement

The authors have declared no competing interest.

## References

1. Moss-Racusin, C.A., et al., Science faculty’s subtle gender biases favor male students. Proceedings of the National Academy of Sciences, 2012. 109(41): p. 16474.

2. Holst, S. and S. Hägg, Positive bias for European men in peer reviewed applications for faculty position at Karolinska Institutet. F1000Res, 2017. 6: p. 2145.

3. Wennerås, C. and A. Wold, Nepotism and sexism in peer-review. Nature, 1997. 387(6631): p. 341–343.

4. Dworkin, J.D., et al., The extent and drivers of gender imbalance in neuroscience reference lists. Nature Neuroscience, 2020. 23(8): p. 918–926.

5. Kaatz, A., et al., Analysis of National Institutes of Health R01 application critiques, impact, and criteria scores: does the sex of the principal investigator make a difference? Academic Medicine, 2016. 91(8): p. 1080–1088.

6. MacNell, L., A. Driscoll, and A.N. Hunt, What’s in a Name: Exposing Gender Bias in Student Ratings of Teaching. Innovative Higher Education, 2015. 40(4): p. 291–303.

7. Astegiano, J., E. Sebastián-González, and C.d.T. Castanho, Unravelling the gender productivity gap in science: a meta-analytical review. Royal Society Open Science, 2019. 6(6): p. 181566.

8. Women, Minorities, and Persons with Disabilities in Science and Engineering. National Center for Science and Engineering Statistics (NCSES) 2019.

9. Malcom, L. and S. Malcom, The double bind: The next generation. Harvard Educational Review, 2011. 81(2): p. 162–172.

10. Sheltzer, J.M. and J.C. Smith, Elite male faculty in the life sciences employ fewer women. Proceedings of the National Academy of Sciences, 2014. 111(28): p. 10107–10112.

11. Pezzoni, M., et al., Gender and the publication output of graduate students: A case study. PLoS One, 2016. 11(1): p. e0145146.

12. Gaule, P. and M. Piacentini, An advisor like me? Advisor gender and post-graduate careers in science. Research Policy, 2018. 47(4): p. 805–813.

13. Hechtman, L.A., et al., NIH funding longevity by gender. Proceedings of the National Academy of Sciences, 2018. 115(31): p. 7943–7948.

14. Cota, R.G., et al., An unsupervised heuristic-based hierarchical method for name disambiguation in bibliographic citations. Journal of the American Society for Information Science and Technology, 2010. 61(9): p. 1853–1870.

15. Song, Y., et al. Efficient topic-based unsupervised name disambiguation. in Proceedings of the 7th ACM/IEEE-CS joint conference on Digital libraries. 2007.

16. Song, Y., et al., Generative models for name disambiguation, in Proceedings of the 16th international conference on World Wide Web. 2007, Association for Computing Machinery: Banff, Alberta, Canada. p. 1163–1164.

17. Hutchins, B.I., et al., Relative Citation Ratio (RCR): A new metric that uses citation rates to measure influence at the article level. PLoS biology, 2016. 14(9): p. e1002541.

18. Lerchenmueller, M.J. and O. Sorenson, The gender gap in early career transitions in the life sciences. Research Policy, 2018. 47(6): p. 1007–1017.

19. Santangelo, G.M., Article-level assessment of influence and translation in biomedical research. Molecular biology of the cell, 2017. 28(11): p. 1401–1408.

20. The SAGE Encyclopedia of Social Science Research Methods. 2004.

21. Hutchins, B.I., et al., Predicting translational progress in biomedical research. PLOS Biology, 2019. 17(10): p. e3000416.

22. Liu, W., et al., Author name disambiguation for PubMed. Journal of the Association for Information Science and Technology, 2014. 65(4): p. 765–781.

23. Song, M., E.H.-J. Kim, and H.J. Kim, Exploring author name disambiguation on PubMed-scale. Journal of Informetrics, 2015. 9(4): p. 924–941.

24. Torvik, V.I. and N.R. Smalheiser, Author name disambiguation in MEDLINE. ACM Transactions on Knowledge Discovery from Data (TKDD), 2009. 3(3): p. 1–29.

25. Kirchmeyer, C., The effects of mentoring on academic careers over time: Testing performance and political perspectives. Human Relations, 2005. 58(5): p. 637–660.

26. Eppstein, D., Fast hierarchical clustering and other applications of dynamic closest pairs. ACM J. Exp. Algorithmics, 2001. 5: p. 1–es.

27. Harris, J.K., et al., Diversify the syllabi: Underrepresentation of female authors in college course readings. PLOS ONE, 2020. 15(10): p. e0239012.

28. Xu, R.F., N.H. Varady, and A.F. Chen, Trends in Gender Disparities in Authorship of Arthroplasty Research. JBJS, 2020. 102(23): p. e131.

