## Supplemental Information for "The effect of mentee and mentor gender on scientific productivity of applicants for NIH training fellowships"

a

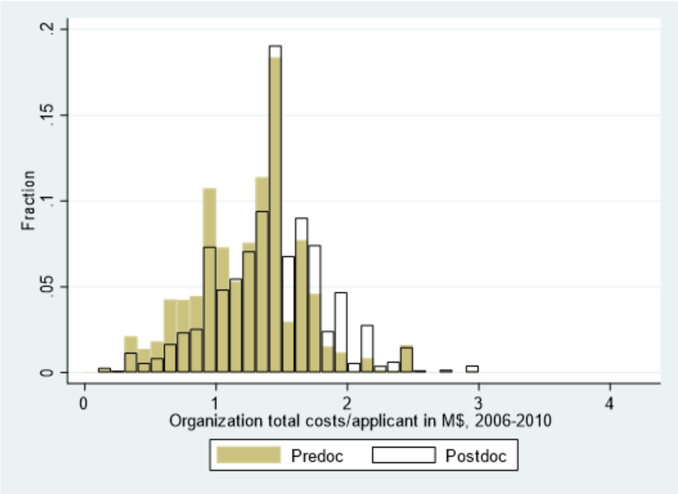


b

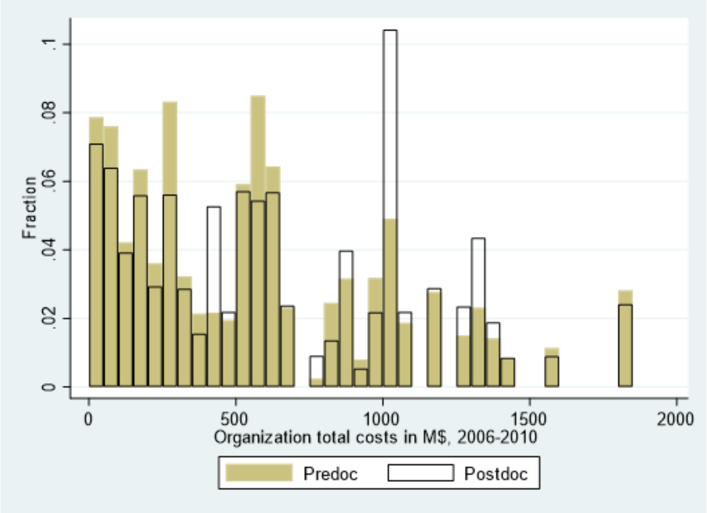


**Figure S1. Distribution of funding to organizations applying for pre- and post-doctoral fellowships.** **(a)** Funding dollars to organizations listed on mentee fellowship applications are analyzed based on each organization's R01 applications in 2006-2010, and presented per applicant. Total costs to the organizations over the entire period of 2006-2010 are presented in units of million dollars, inflation adjusted to 2019$ using BRDPI, and then divided by the number of unique Principal Investigator (PI) applicants for R01s from that organization in 2006-2010. This provides the total costs awarded per organizational PI applicant, taking into account organizations that have a relatively small number of PIs but high total costs per PI. **(b)** Same data as in (a) but presented as total costs to the organizations, rather than as a per applicant metric. For both (a) and (b), the fractions are presented within each career stage category, i.e. all clear bars (organizations from mentee post-doctoral fellowship applications) sum to 1; same for brown bars (organizations from mentee pre-doctoral fellowship applications).

a

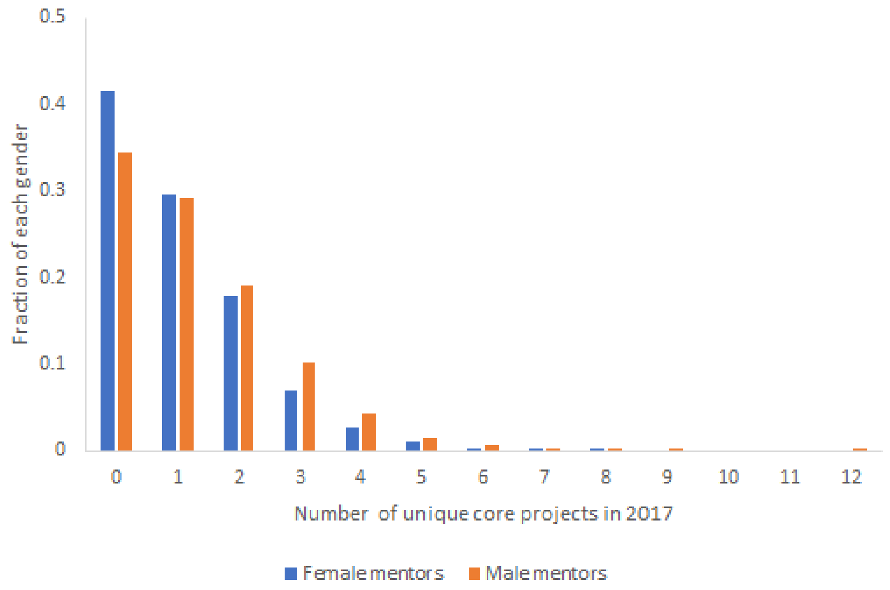


b

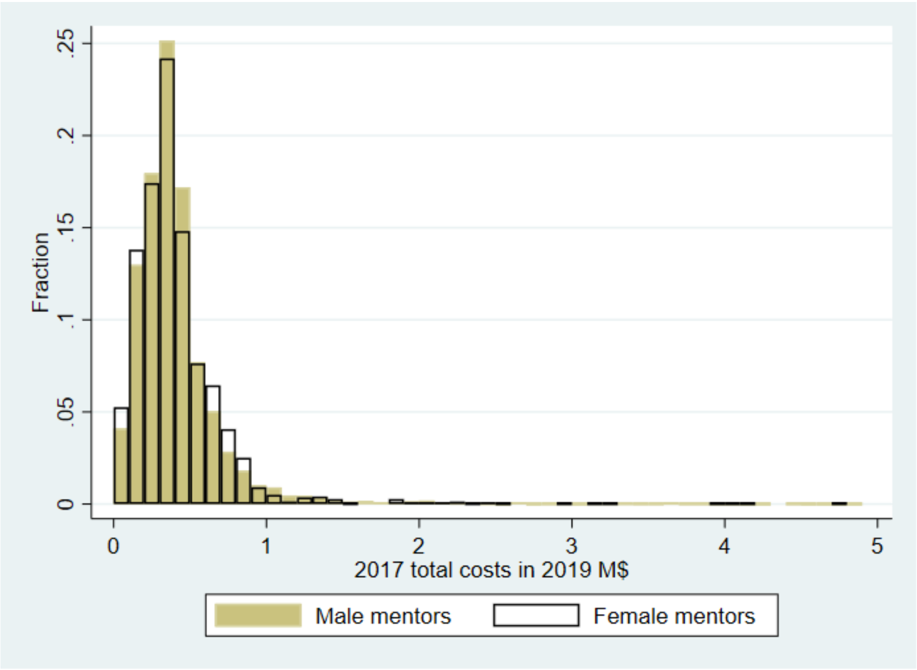


**Figure S2. Distribution of mentor award number and award size, by gender.**
In order to understand the effect caused by normalizing to mentor dollar resources (Figure 5), mentor award numbers and sizes were further analyzed by gender. **(a)** The fraction of mentors with the given number of unique awarded projects in 2017. The fractions are presented within each gender, i.e. all the blue bars (female mentors) sum to 1; same for the orange bars (male mentors). Using a Mann-Whitney test, the number of awards for male mentors is significantly higher than for female mentors (p-value<0.0001). **(b)** The size of each award received by mentors by mentor gender in 2017. The unit of analysis is the award-mentor with each entry representing the estimated total costs allocated to the

given mentor-PI. The total costs are in units of million dollars, inflation adjusted to 2019$ using BRDPI. The fractions are presented within each gender, i.e. all clear bars (female mentors) sum to 1; same for brown bars (male mentors). PIs with no funding in 2017 are not included in this figure. Awards above $5 million have also been excluded. Using a Mann Whitney test, the distributions of award sizes are not significantly different by mentor gender.
